# Relationship between swimming velocity and trunk twist motion in short-distance crawl swimming

**DOI:** 10.1101/2022.09.04.506549

**Authors:** Hiroki Hyodo, Daiki Koga, Yasuo Sengoku, Tadashi Wada

**Author notes:** Corresponding author (HH).

## Abstract

This study aimed to estimate the trunk twist angle from the shoulder and hip rotation angles in short-distance crawl swimming and to elucidate the twist motion of the relationship between the trunk and the rotation angular velocity in response to changes in swimming speed. Swimming speed during the experimental trials was computed from the subject’s best times in the 50 and 100m crawl swims. Wireless self-luminous LED markers were attached to seven locations on the body. The actual coordinate values of the LED markers were obtained using 18 cameras for underwater movements and 4 on the water for above-water movements. A comparison of the rate of change between trials revealed a high correlation (r = 0.722, p < 0.01) between the twist angle and shoulder rotation angular velocity in the push phase. In the same phase, a high correlation (r =0.748, p < 0.01) was also found between the twist angle and the angular velocity of hip rotation. These results suggest that swimmers increase the twist angle of their trunks to obtain a higher swimming speed. Moreover, the trunk muscle group increases its activity before starting the main motion of the twist back motion, and stretch-shortening cycle (SSC) motion may be performed in the trunk.

## Introduction

A swimming race is a competition in which swimmers travel a defined distance from the start to the finish line in a defined discipline and within the predefined time required to reach the finish line. The race is divided into the start phase (i.e., 15 m from the start), the turn phase (i.e., 5 m before the turn), the finishing phase (i.e., 5 m before the finish), and the stroke phase (i.e., phases other than start, turn, and finish). The stroke phase is the most common, and it is important to obtain a high swimming speed by upper limb strokes to improve performance (Wakayoshi, 1992). There are four types of swimming strokes in competitive swimming: butterfly, backstroke, breaststroke, and crawl. Crawl swimming is considered the swimming stroke that can attain the highest swimming speed, and requires the swimmers to maintain a horizontal position in the water and exert propulsive force in an environment without a support point (Hollander et al., 1986).

Regarding the relationship between swimming speed and swimming motion, Matsuda et al. (1998) reported that the characteristics of swimming change with changes in stroke frequency as swimming speed changes. The average swimming speeds in the 50-m crawl and 100-m crawl races, which are both short-distance swimming events in competitive swimming, were compared to world records: the world record for the 50-m crawl was 20.91 s, with an average swimming speed of 2.39 m/s, and the world record for the 100 m crawl was 46.91 s, with an average swimming speed of 2.13 m/s, a difference of 0.26 m/s. Similarly, in short-distance events at the national competition level, the difference in average swimming speed is more than 2.0 m/s. Narita et al. (2018) reported that the body receives from the fluid while swimming increases in proportion to the third power of the swimming speed, suggesting that higher swimming speed environments may produce greater resistance. These results suggest that the swimming characteristics of the 50-m crawl swimmer and the 100-m crawl swimmer may differ because of the difference in swimming speed. Therefore, it is necessary to distinguish between 50-m crawl swimming and 100-m crawl swimming in terms of the characteristics of a swimming motion during a race. Focusing and by focusing on the changes in a swimming motion is expected to discover findings that improve maximum swimming speed in short-distance events. In crawl swimming, it is known that the propulsive force obtained by upper limb motion is greater than that of other swimming strokes. It has been reported that upper limb motion accounts for approximately 90% of the total whole-body propulsive force (Watkins et al.,1983; Hollander et al., 1986; Deschodt et al., 1999; Gourgoulis et al.)

Furthermore, crawl swimming involves a rotational motion of the trunk around the long axis in conjunction with upper limb motion, and this motion is defined as a trunk rotation (Yanai, 2003). In crawl swimming, it has been reported that trunk rotation affects the direction and speed of internal and external hand motions during the pull phase, which is the phase before the hand enters the water and moves just below the shoulder (Grimston & Hay, 1986; Payton et al., 2002). In a previous study that measured the fluid force generated at hand using pressure distribution measurements at the swimmer’s hand, it was also reported that an increase in hand velocity contributes to an increase in the propulsive force exerted at hand because the fluid force increases in proportion to the square of the velocity (Narita et al., 2017; Tsunokawa et al., 2019; Tsunokawa et al., 2019).Furthermore, regarding the relationship between hand motion and trunk rotation, it has been confirmed that an increase in shoulder rotation angular velocity affects the vertical hand velocity in the push phase. An increase in vertical hand velocity in the push phase contributes to the lift exerted by the hand and increases hand propulsion lift (Kudo et al.,2017; Schleihauf, 1983). These results indicate that an increase in the angular velocity of the rotation at the trunk during the swimming motion is important to improve the propulsive force. It is necessary to elucidate the factors that cause the increase in angular velocity to improve performance. Hyodo et al. (2021) investigated shoulder and hip rotation changes at different swimming speeds during short-distance races. The results showed that the hips’ rotation angle peak changed earlier at higher swimming speeds during the phase time until the peak of the rotation angle in the pull phase. In addition, the phase time of the shoulder and hip rotation changed. At higher swimming speeds, the waist shifted from a negative direction toward the pool bottom (ventral side) to a positive direction toward the water surface (dorsal side) earlier. As a result, the authors report a difference in phase time between the shoulder and hip rotation and possibly increasing suggests the trunk twisting motion of the trunk may become larger. Therefore, clarification of the twisting motion of the trunk may contribute to elucidating the increase in the angular velocity of rotation, which has a significant effect on swimming performance.

In a previous study on the trunk twisting motion, Wada et al. (2003) conducted a three-dimensional motion analysis during volleyball spike motion. They calculated the trunk twist angle based on the difference between the shoulder and waist rotation angle. They found a positive correlation between the trunk twist angle and the shoulder rotation angular velocity during a volleyball spike action. The same study also reported that the angular velocity of shoulder rotation during the roll back phase of the spike motion affects hand velocity. In crawl swimming, hand speed also affects performance. It is possible that hand velocity, which is important for improving swimming speed, is affected by twisting motion. However, no studies focus on the relationship between the angular velocity of trunk rotation and trunk twisting motion in crawling swimmers. Furthermore, the parameters in these studies were only between subjects, and no previous studies have focused on changes in swimming motion at different speeds.

The purpose of this study was to analyze three-dimensional motion during crawl swimming using a motion capture system, to calculate the trunk twist angle from the shoulder and hip rotation angles during short-distance crawl swimming, and to clarify the relationship between the trunk twisting motion and the rotation angular velocity in response to changes in swimming speed.

## Materials and methods

### Analyte

Ten subjects (Table 1) participated in this study (age: 21.0 ± 2.5 years, height: 179.8 ± 6.7 cm, weight: 77.3 ± 5.9 kg). All subjects trained six days a week and all had participated in national competitions.

If the subjects were minors, their guardians were informed prior to the experiment that they could withdraw their consent to participate in this study of their own free will at any time. Even after informed consent, they would not be treated unfavorably because of such withdrawal. In addition, we informed them that personal information obtained during the research would be handled appropriately to prevent leakage or loss. The purpose and methods of this study were explained to the subjects, and their informed consent to participate in the experiment was obtained.

**Table 1.** Physical Characteristic Performance Level of Subjects.

| Subject | Age<br>(years) | Height<br>(cm) | Weight<br>(kg) | Best Record of<br>50 m FreeStyle<br>(s") | Fina Point | Best Record of<br>100 m FreeStyle<br>(s") | Fina Point |
| --- | --- | --- | --- | --- | --- | --- | --- |
| A | 21 | 189 | 80 | 23.37 | 716.3 | 51.74 | 745.3 |
| B | 20 | 169 | 71 | 23.94 | 666.3 | 53.04 | 691.8 |
| C | 23 | 180 | 83 | 23.41 | 712.6 | 51.34 | 762.8 |
| D | 18 | 180 | 76 | 24.06 | 656.4 | 51.72 | 746.1 |
| E | 22 | 180 | 74 | 23.59 | 696.4 | 51.91 | 738.0 |
| F | 18 | 185 | 72 | 23.40 | 713.5 | 52.11 | 729.5 |
| G | 21 | 169 | 72 | 23.87 | 672.2 | 50.95 | 780.5 |
| H | 20 | 183 | 75 | 24.26 | 640.3 | 51.43 | 758.8 |
| I | 20 | 175 | 79 | 23.95 | 665.5 | 51.33 | 763.3 |
| J | 27 | 188 | 91 | 21.67 | 898.4 | 48.35 | 913.3 |
| mean | 21.0 | 179.8 | 77.3 | 23.55 |  | 51.39 |  |
| SD | 2.5 | 6.7 | 5.9 | 0.69 |  | 1.15 |  |

### Measurement method

The experiment was conducted in a reflux water tank (Igarashi Industries, Japan), where swimming speed could be controlled by artificially flowing water inside the pool. The sides and bottom of the observation area were covered with glass. Swimming speed during the experimental trials was calculated from the subject’s best times in the 50- and 100- m crawl swims. To eliminate the effect of diving, the starting phase was defined as the period from the start signal until the swimmer reached the 15- m point. The average swimming speed of the phase was used, excluding the starting phase from each best time as the set swimming speed. After warming up sufficiently, the participants performed a100-m crawl average swimming speed test (“V100 m”) and a 50-m crawl average swimming speed test (“V50 m”) for 10 s each in the first and second sessions, respectively. To prevent changes in swimming speed during the test, markers were placed at the bottom of the pool so that the subject did not move from the measurement section. The subject’skicking motion was the same as that in a race, i.e., a six-beat kick with six kicking motions in one stroke. The test was performed without breathing to eliminate the effect of breathing on the stroke motion.

### Phasing of upper limb strokes

In this study, upper limb stroke motion was divided into phases (Fig 1) based on the method by Chollet et al. (2000) to analyze the relationship between each variable and the swimming motion. The phases of the swimming motion were divided using an absolute coordinate system rather than a moving coordinate system fixed to the swimmer. A cycle of strokes was divided into four phases as follows. The glide phase is the period between the point where the Z-coordinate of the fifth metacarpophalangeal joint becomes negative (i.e., water entry point) and the point where the Y-coordinate of the fifth metacarpophalangeal joint begins to move in the opposite direction of the swimmer’s swimming direction when the Z-coordinate of the fifth metacarpophalangeal joint is 0 at the water surface. The pull phase was defined as the period from the end of the glide phase until the Y-coordinate of the fifth metacarpophalangeal joint, reaching just below the Y-coordinate of the acromion. The recovery phase was defined as the period from the end of the push phase to the start of the glide phase (the end of one cycle).

**Figure 1.**
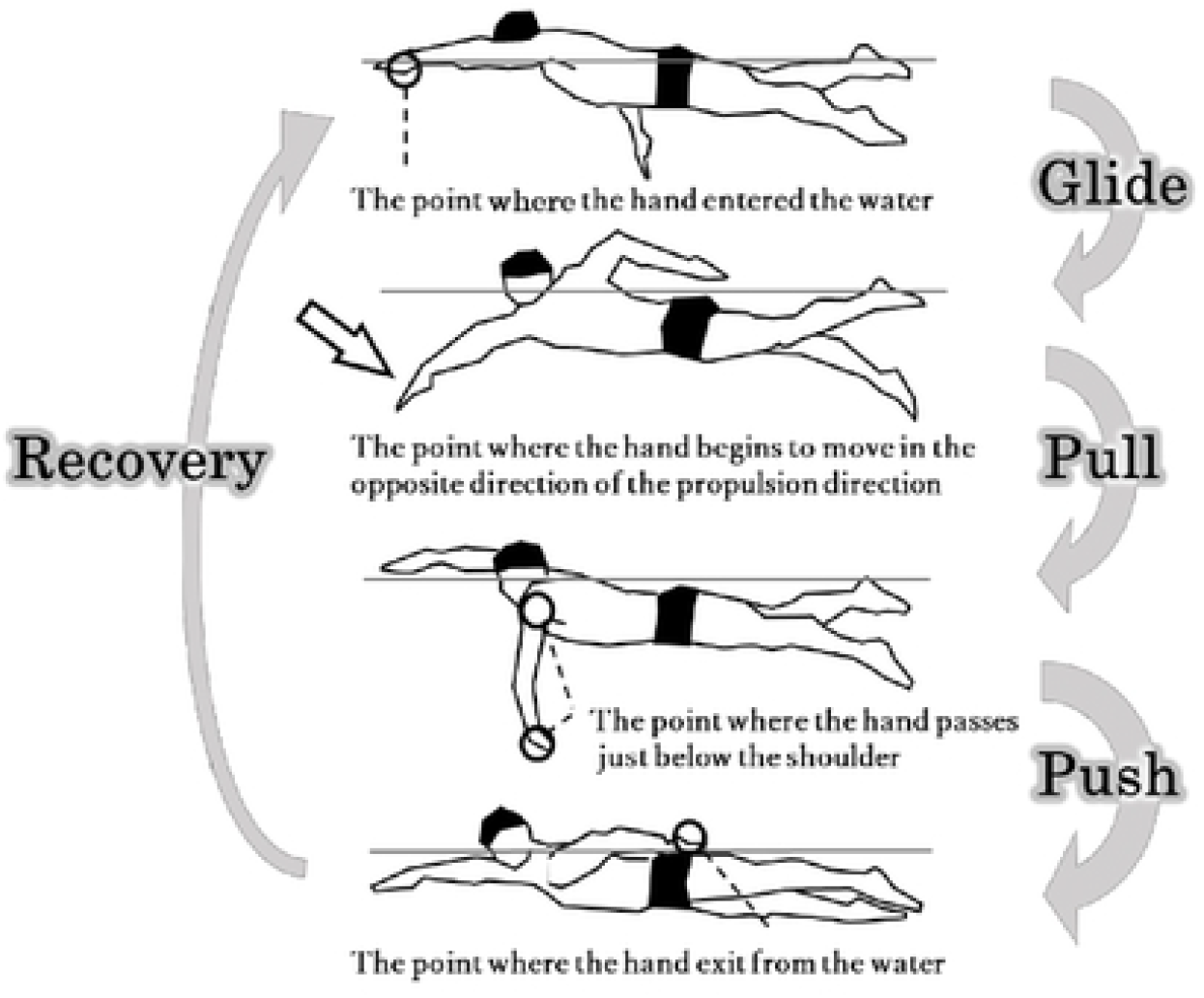
Phase Division of Upper Limb Stroke.

### Three-dimensional motion analysis and measurement items

To investigate variables such as upper limb and trunk motion of the subject, seven locations on the body (i.e., top of the head, right and left acromions, right and left fifth metacarpophalangeal joints, and right and left greater trochanters) were marked using wireless self-luminous LED markers (“Kirameki”, Nobby Tech Inc., Japan), referring to the method of Hyodo et al. (2021) (Fig 2). Motion capture cameras (Opti Track, Acuity Inc., Japan) were installed around the experimental turning basin. To measure underwater motion, five cameras were set up on each side of the swimmer and eight on the lower side (for a total of 18 cameras used to capture underwater motion. Four cameras were set up above the turning basin for the above-water motion to acquire real coordinate values for LED markers in the calibration volume. The cameras were positioned at a sampling frequency of 100 Hz. The angle of view was adjusted so that the markers attached to the body could be photographed from two or more cameras (Fig 3). To avoid the misidentification of the LED markers due to reflections from the water and glass surfaces during the test and calibration, the room was kept dark with curtains to block sunlight from the outdoors.

**Figure 2.**
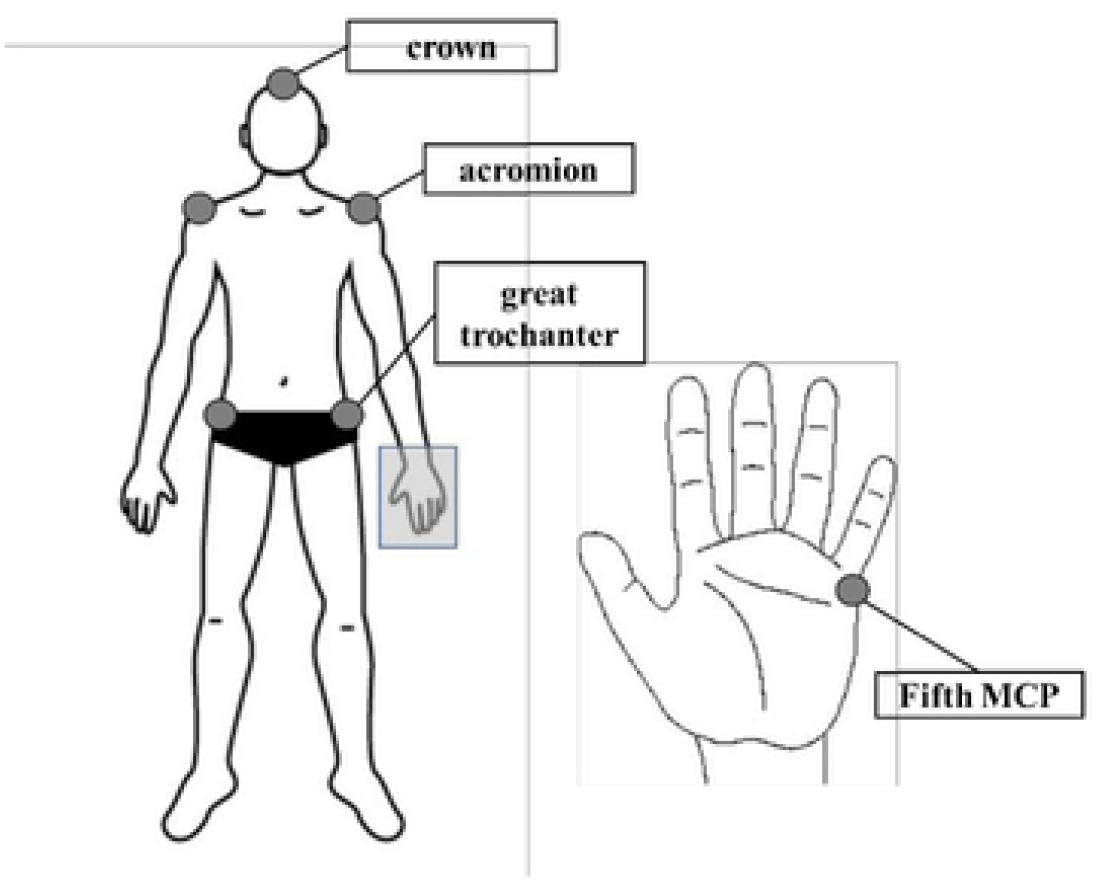
Diagram of Marking Points.

**Figure 3.**
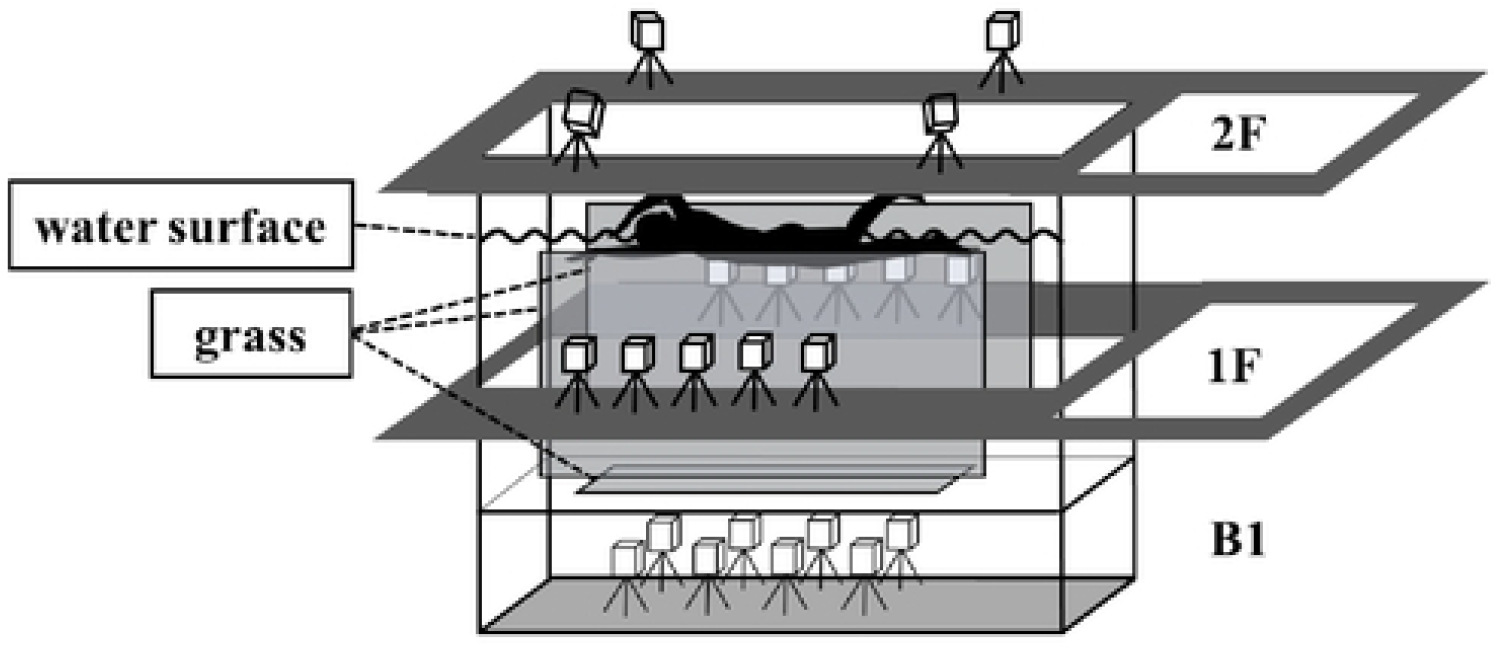
Diagram of the Experimental Setting.

For the analysis, image analysis software (VENUS 3D R, Nobby Tech Inc., Japan) was used to define a virtual perpendicular line to the space from the coordinate values (X, Y)on the two-dimensional plane using the angle and distance of the two-dimensional plane obtained from each camera with an epipolar matching algorithm. The 3D space was constructed from the 3D coordinate values of the body as follows. A fixed right-handed coordinate system was used, with the subject’s left and right directions on the X-axis (i.e., positive direction on the left side), the direction of motion on the Y-axis (i.e., positive direction in the direction of propulsion), and the vertical direction on the Z-axis (i.e., positive direction on the parietal side) (Fig 4). The obtained real coordinates were smoothed using a low-pass filter to reduce noise by omitting frequencies above the outliers.

**Figure 4.**
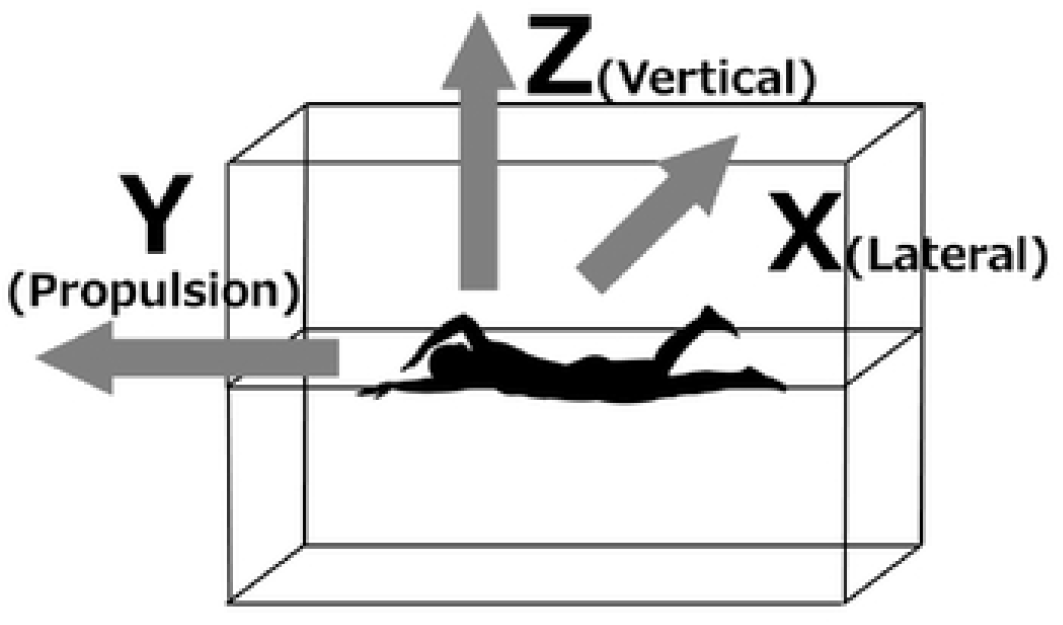
Coordinate Axis Definition.

Before the experiment, more than 1,500 samples were acquired with each camera by waving the wand within the angle of view of each camera to perform dynamic calibration using a dedicated calibration wand to construct a three-dimensional coordinate space for the measurement range. Dynamic calibration was performed separately underwater and above water to eliminate light refraction from the water surface. After calibration, the standard error of motion capture was less than 0.5 mm in and on the water. Then, a calibration LED marker plate with three points arranged in an L-shape was floated on the water’s surface for a 30-s calibration period to combine the underwater and above-water three-dimensional coordinate spaces. The water surface was identified by photographing the marker positions from underwater and above water, and the origin of the space was determined by aligning the coordinate axes of underwater and above water. Because the plate has a thickness and there is a gap between the markers placed in the water and on the water surface, the length of the gap was measured and reflected to fill in the difference when aligning the coordinate axes. In addition, the calculation error was calculated by measuring the length of the wand of a fixed size at multiple locations within the angle of view after calibration (Koga et al., 2021). One hundred samples of wand lengths were measured at each of three locations within the measurement range (front, middle, and rear), and the maximum error was less than 0.5 mm.

The analysis elements of this study are as follows. The abbreviations of each item are shown below. Stroke frequency (strokes/s), hereafter abbreviated as SF (stroke frequency).For SF, one cycle was defined as the stroke from the direction of the position where the Z-axis of the hand became negative to the position where it became positive and then became negative again. Stroke length (m/stroke), hereafter abbreviated as “SL (stroke length),” was calculated. SL was calculated by dividing SF by the set swimming speed, referring to the method of Kennedy et al. (1990).

The shoulder rotation angle (degrees), hereafter abbreviated as “ShR (shoulder roll),”was defined as the angle between the horizontal plane and the straight line connecting the right and left acromions when projected onto the XZ plane. The rotation angle was defined as 0 ° when the swimmer’s shoulders were parallel to the horizontal plane. The negative rotation angle was defined as the value in the direction in which the right shoulder was down. The hip rotation angle (degrees), hereafter abbreviated as “HiR (hip roll),” was defined as the angle between the horizontal plane and the straight line connecting the left and right obliques projected onto the XZ plane. The rotation angle was defined as 0 ° when the swimmer’s hips were parallel to the horizontal plane. The negative rotation angle was defined as the value in the direction in which the right hip was lowered. Trunk twist angle (degrees), hereafter abbreviated as “TA (twist angle).” TA was calculated from the difference between ShR and HiR, referring to the method of Wada et al. (2003) (Fig 5). The timing when the angle difference showed the maximum value was defined as the maximum twisting motion, and the value at this time was defined as TA. Shoulder rotation angular velocity (rad/s) is abbreviated as ShRAV (shoulder roll angular velocity). Hip rotation angular velocity (rad/s), hereafter abbreviated as HiRAV (hip roll angular velocity). With time, the angular velocities of shoulder and hip rotations were calculated by differentiating the displacements of ShR and HiR. Because the direction of motion switches from negative to positive during the pull phase, we defined the phase in which the right shoulder moves toward the pool bottom and rotates in the negative direction as the pull roll phase, the phase in which the shoulder moves toward the water and rotates in the positive direction as the pull roll back phase, and the phase from the end of the pull phase to hand ejection as the push phase, and calculated the average angular velocity during each phase.

**Figure 5.**
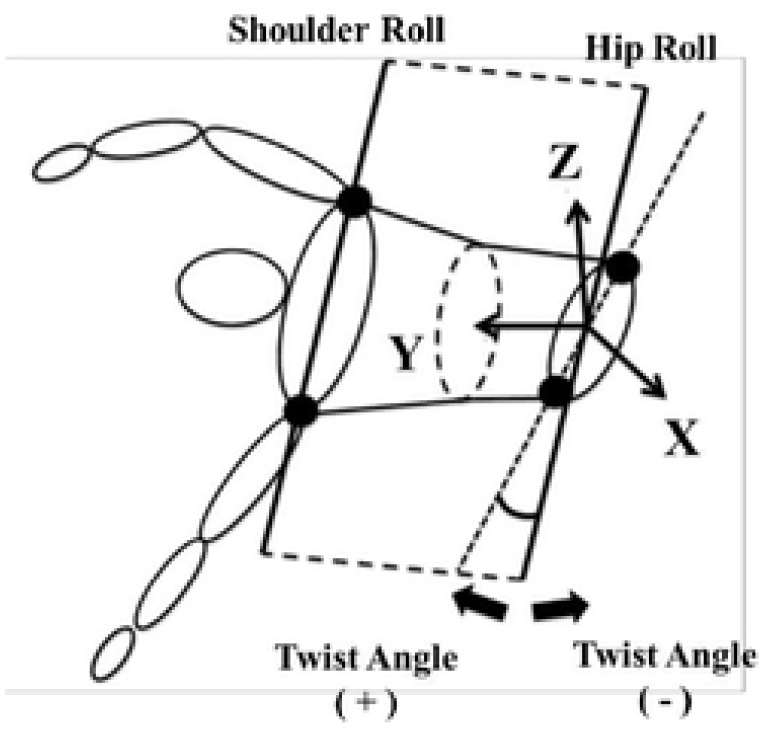
Definition of Trunk Twist Angle.

The timing of the maximum shoulder and hip rotation angles was analyzed, abbreviated as “peak time.” The relative time (%) from the beginning of the phase to the maximum rotation angle is shown for the phases in which the maximum rotation angle was observed. The start of the phase was set at 0%, and the end of the phase was set at 100%. The timing at which the shoulder rotation angle reached its maximum value (ShR peak time) and that at which the hip rotation angle reached its maximum value (HiR peak time) were calculated. To make comparisons at different swimming speeds, the time taken for the target phase of each trial was normalized to 100%. The time from the beginning of the phase to the maximum value of the rotation angle was expressed as phase percentage (%).

### Statistical analysis

The Shapiro–Wilk test confirmed the normality of the data. Significant differences between trials when the swimming speed was varied were analyzed using the corresponding t-test. Correlations between variables were analyzed by performing an uncorrelated test using Pearson’s correlation coefficient. Following Hopkins, Marshall, Batterhan, and Hanin (2009), correlation coefficient thresholds were defined as 0.1, 0.3, 0.5, 0.7, and 0.9 for small, moderate, large, very large, and near-perfect, respectively. All statistical analyses were performed using SPSS Statistics ver. 25 (IBM, USA), with a risk rate of less than 5 % as the criterion for statistical significance.

## Result

### Set swim speed (SV), SF, and SL for each trial

Table 2 shows each trial’s mean and standard deviation of the set swimming velocity (SV),SF, and SL. The mean swimming speeds calculated from the best time excluding the start phase, were 1.97 ± 0.06 m/s for V50 m and 1.87 ± 0.04 m/s for V100 m, with V50 m having a significantly higher value (t = 7.21, p < 0.01). SF was 63.08 ± 3.93 strokes/min for V50 m and 56.69 ± 3.26 strokes/min for V100 m, with V50 m showing significantly higher values (t = 7.29, p < 0.01). SL was 1.88 ± 0.14 m/stroke for V50 m and 1.98 ± 0.11 m/stroke for V100 m, with V50 m showing significantly lower values (t = 4.25, p < 0.01).

**Table 2.** Mean Values of Each Variable at V50 m and V100 m.

| Subject | SV (m / s) |  | SF (stroke / min) |  | SL (m / stroke) |  |
| --- | --- | --- | --- | --- | --- | --- |
|  | V50 m | V100 m | V50 m | V100 m | V50 m | V100 m |
| A | 1.97 | 1.85 | 65.22 | 58.82 | 1.81 | 1.89 |
| B | 1.94 | 1.81 | 62.50 | 53.57 | 1.86 | 2.03 |
| C | 1.97 | 1.86 | 59.41 | 54.55 | 1.99 | 2.05 |
| D | 1.92 | 1.86 | 65.22 | 59.41 | 1.77 | 1.88 |
| E | 1.98 | 1.85 | 67.42 | 58.25 | 1.76 | 1.91 |
| F | 1.99 | 1.84 | 61.86 | 51.72 | 1.93 | 2.13 |
| G | 1.93 | 1.88 | 69.77 | 60.61 | 1.66 | 1.86 |
| H | 1.90 | 1.86 | 55.56 | 52.63 | 2.05 | 2.12 |
| I | 1.94 | 1.88 | 63.83 | 61.22 | 1.82 | 1.84 |
| J | 2.14 | 1.98 | 60.00 | 56.07 | 2.14 | 2.12 |
| mean | 1.97 ** | 1.87 | 63.08 ** | 56.69 | 1.88 ** | 1.98 |
| SD | 0.06 | 0.04 | 3.93 | 3.26 | 0.14 | 0.11 |

Significant difference between conditions. \* $p < 0.05$ , \*\* $p < 0.01$

### Comparison of different swimming speeds

Table 3 shows the mean and standard deviation of the peak time for ShR, HiR, TA, ShRAV, HiRAV, and each rotation. For ShR, the values were 52.9° ± 12.9° for V50 m and 55.8° ±14.0° for V100 m, with significantly lower values for V50 m (t = 2.22, p < 0.05). For HiR, there were no significant differences between trials. For TA, the values were 34.0° ± 7.2°for V50 m and 30.6° ± 8.9° for V100 m, with significantly lower values for V50 m (t =2.75, p < 0.05). There were no significant differences between trials in all phases for the ShRAV and HiRAV variables for angular velocity. Because ShR and HiR showed their maximum values during the pull phase, the time until ShR and HiR showed their maximum values during the pull phase was calculated. The results showed that V50 m was significantly lower than V100 m trials at HiR peak time (t = 4.45, p < 0.01).

**Table 3.**
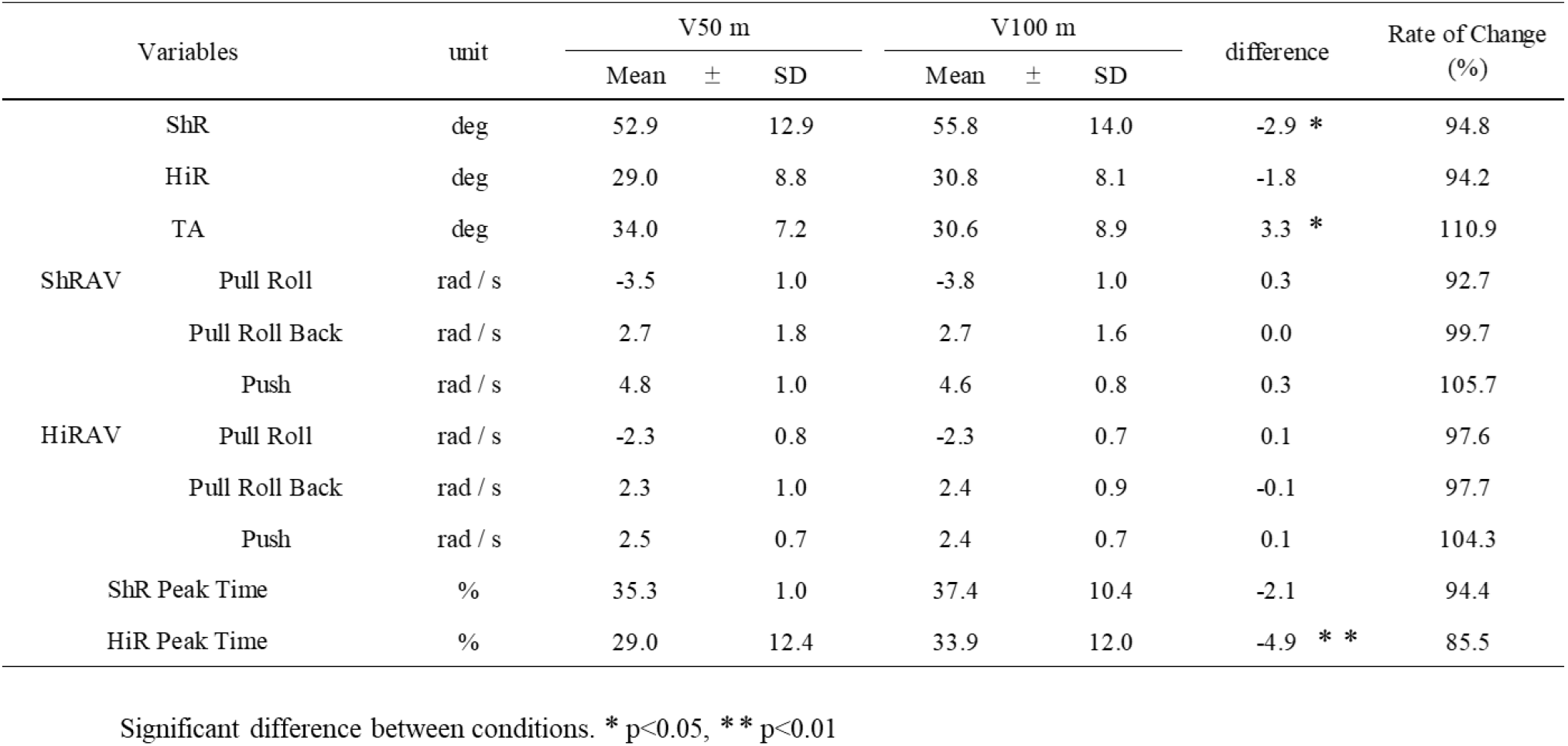
Mean Values of ShR, HiR and TA Variables at V50 m and V100 m.

### Relationship between rotation angular velocity and TA

Table 4 shows the correlation coefficients between the percent change in TA at V50 m relative to V100 m and the percent change in ShRAV and HiRAV for each phase. The results of the correlation analysis showed that there was no correlation between the shoulders and hips in the pull roll phase. A moderate correlation (r = 0.575, p < 0.05) was found between TA and the rate of change of ShRAV during the pull roll back phase. In the push phase, a high correlation (r = 0.722, p < 0.01) was found between TA and the rate of change in ShRAV, and a high correlation (r = 0.748, p < 0.01) between TA and the rate of change in HiRAV. Fig 6 shows the correlation diagram between TA and the rate of change of ShRAV during the push phase, where the correlation coefficient was particularly high. Fig 7 shows the correlation diagram between TA and HiRAV.

**Table 4.**
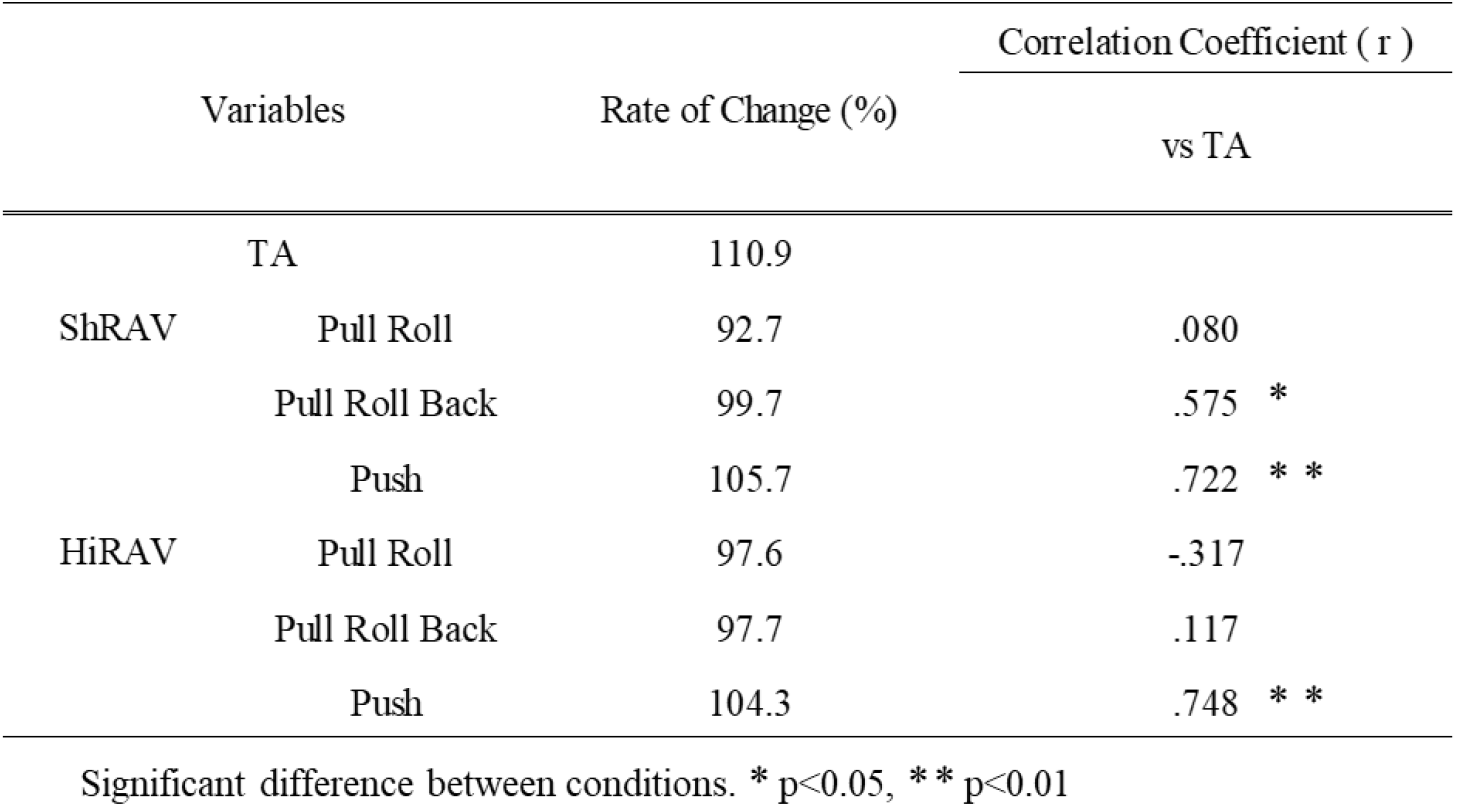
Correlation Coefficients between the Percent Change in TA at V50 m Relative to V100 m and the Percent Change in ShRAV and HiRAV for Each Phase.

**Figure 6.**
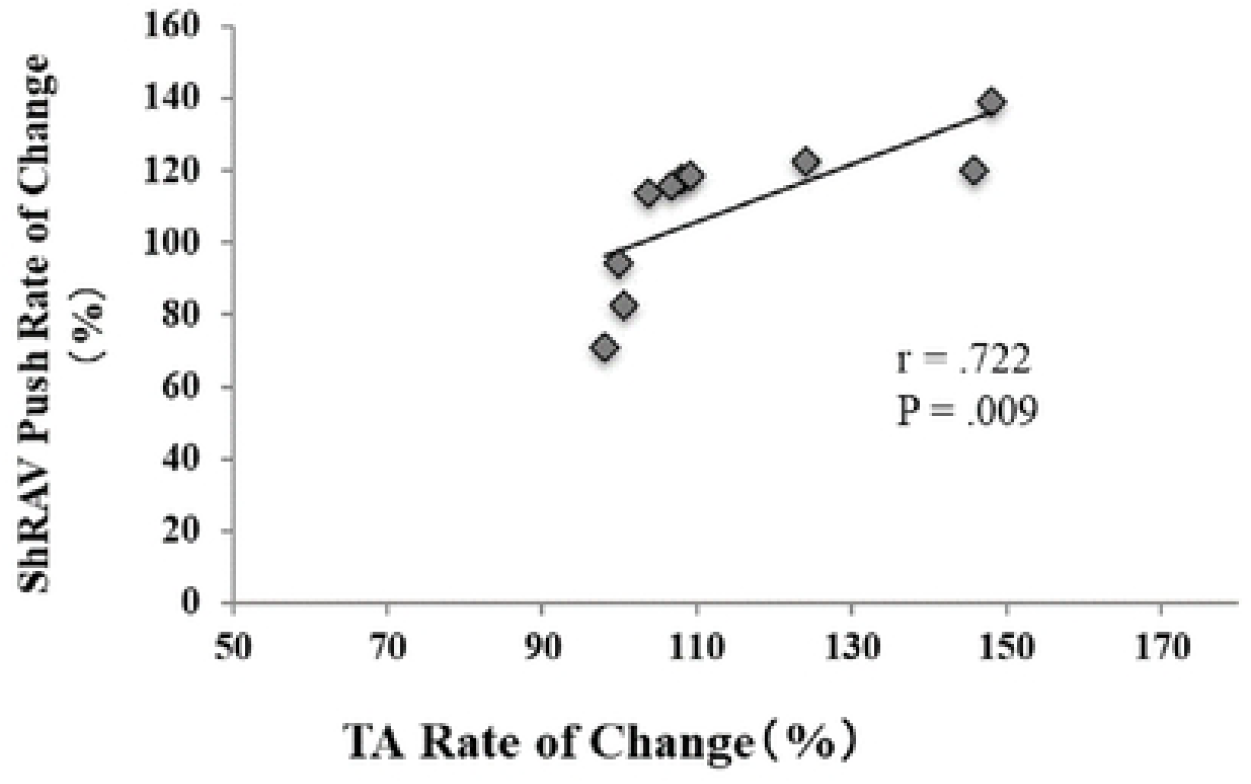
Correlation Diagram between TA and the Rate of Change of ShRAV during the Push Phase.

**Figure 7.**
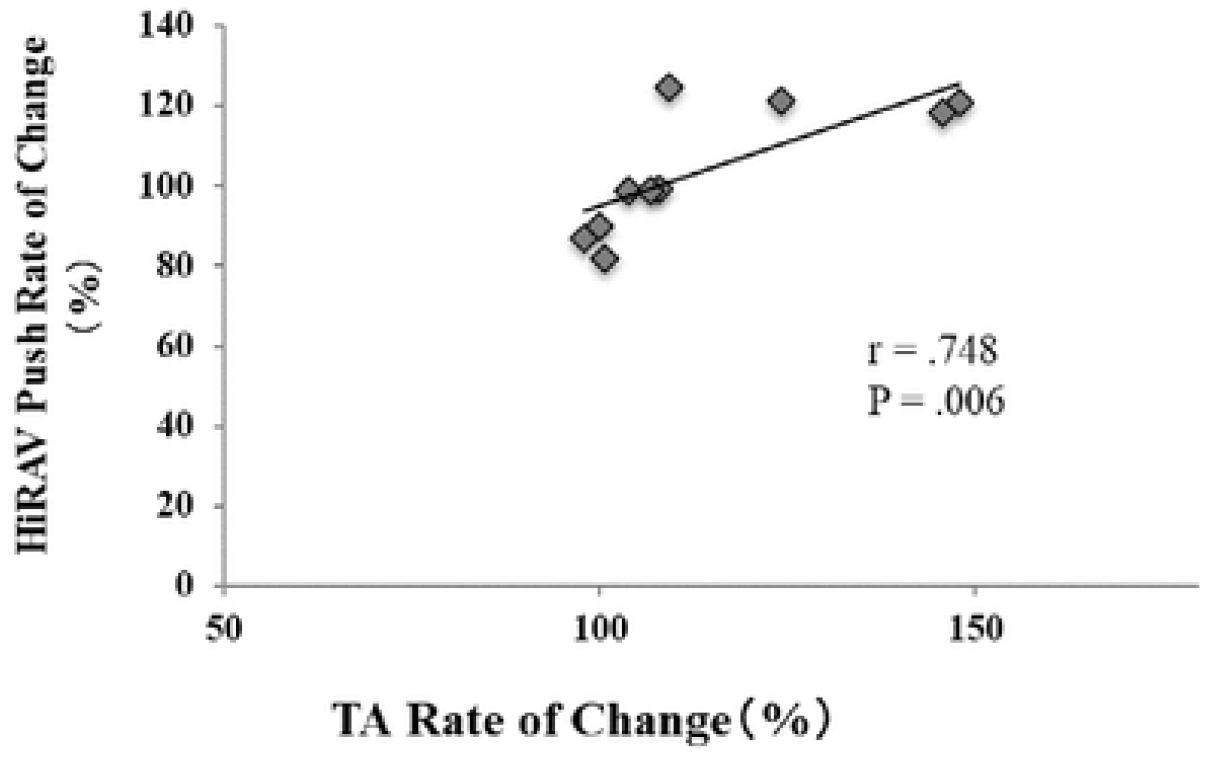
Correlation Diagram between TA and the Rate of Change of HiRAV during the Push Phase.

## Discussion

### Comparison of ShR, HiR, and TA between trials

ShR for each trial was significantly lower for V50 m than for V100 m. No significant differences in HiR were observed between the trials. In a previous study, Yanai (2003) reported an association between ShR and SF, with ShR showing a decrease with increasing SF. Although the set swimming speed in the present study was higher than that in the previous studies because the set swimming speed was tested in the short-distance events, SF was significantly higher in the V50 m and the ShR was lower, indicating that the results were similar in the short-distance events as in the previous studies of the set swimming speed in the middle and long-distance events.

TA calculated from the difference between ShR and HiR showed significantly higher values for V50 m than for V100 m. Hyodo et al. (2021) reported that the time at which the trunk motion angle showed a maximum value was significantly lower with increasing swimming speed at the phase time until the HiR showed a maximum value. The same results were shown in the present study, suggesting that the early shift of only the hip rotation motion from the negative to the positive direction of the rotation motion increased the trunk twist angle by increasing the time for the hip rotation motion to turn and rotate in the positive direction and the shoulder rotation motion to rotate in the negative direction, with each rotating in the opposite direction. Furthermore, Welch et al. (1995) investigated the trunk twisting motion during the batting motion and confirmed that the rotation of the hips prior to that of the shoulders increases the activity of the main active muscles such as the external obliques and the latissimus dorsi before the main motion of the shoulders begins, by performing the turnover motion earlier, and by increasing the trunk. The results confirmed that the trunk muscle group’s stretch-shortening cycle (SSC) motion was performed. This has been reported to increase the moment exerted on the shoulder, thereby increasing the angular velocity. Therefore, this study analyzed the angular velocities of shoulder and hip rotation.

### Comparison of rotation angular velocity between trials

Although there was no significant difference in ShRAV between the trials in all phases, the angular velocity change from negative to positive direction was observed in the pull phase of both trials for the rotation turnover. ShRAV was then highest during the push phase. Kudo et al. (2017) reported that the angular velocity of shoulder rotation increased from the second half of the pull phase, with maximum values during the push phase. This study obtained similar results for short-distance crawl swimming race speed. Furthermore, in a study conducted by Kudo (2017) in a hydrostatic environment on the hydrodynamic forces generated by hand, he reported that the shoulder rotation angular velocity increased in the push phase increases the hydrodynamic forces at the velocity of the hand by affecting the hand in the same phase, thus contributing to the propulsive lift generated in the push phase. The same results were obtained in the experiment conducted in this study. However, the test was conducted in a reflux water tank to control the swimming speed. Therefore, the propulsive force may increase the shoulder rotation angular velocity in the static water environment and a reflux water tank in the push phase.

No significant differences were found between trials in HiRAV (Psycharakis, 2008). He investigated the relationship between the shoulder and hip rotation and the upper and lower extremities from their respective frequencies. The results showed that the rotation of the hips was greatly influenced by the kicking motion, which was performed with the lower limbs, and showed that the direct relationship between the rotation of the hips and the hand motion was small. Furthermore, in this study, the waist rotation was indirectly related to ShRAV by changing the timing in the pull phase as swimming speed increased, contributing to an increase in TA. These results inferred that hip rotation motion did not directly contribute to hand propulsion but might have affected SF and hand propulsion by influencing ShRAV. Therefore, in the future, a more detailed investigation is needed on the relationship between hip rotation and kicking motion, which is considered highly related to hip rotation motion.

### Relationship between the rate of change of TA and the rate of change of rotation angular velocity between trials

The rate of change between trials at V 50 m and V 100 m was calculated to compare the change in swimming speed between trials. The correlation between the rate of change in TA and the rate of change in ShRAV and HiRAV in each phase was analyzed. For ShRAV, significant correlations were found between the rate of change in TA and the change in the pull-roll-back phase and the rate of change in the push phase (Fig 6). In HiRAV, a significant correlation was found between the rate of change in TA and the rate of change in the push phase (Fig 7). Takahashi et al. (2018) reported that for the trunk twisting motion, increasing the activity of the external obliques and the latissimus dorsi muscles, which are the primary muscles of the trunk twisting motion, before the start of the main motion, increases the moment of the shoulder exerted. A previous study by Hiruma et al. (2010) also reported that the trunk twisting motion was accompanied by the SSC motion of the trunk muscle groups, which contributed to an increase in ShRAV of the twist back after ShR reached its maximum value. This study also showed a high correlation between the change in TA and the rate of change in the pull roll back phase and the push phase, which are the twist back phase in ShRAV, similar to previous studies that investigated the trunk twisting motion in other sports. Therefore, it was inferred that an increase in TA during short-distance crawling induced the SSC motion of the trunk muscle groups, which affected ShRAV during the twist back phase (i.e., pull-roll-back phase and push phase) after ShR reached its maximum value.

Furthermore, these results were related to hand velocity during the push phase, which is considered to have a significant effect on the propulsive force during short-distance crawl swimming, as reported by Kudo et al. (2017) and Koga et al. (2020). The increase in TA is a very important index for improving the propulsive force. However, Maglischo (1993) reported that the rotation motion during crawling might be caused by the buoyancy moment generated by the shoulder sinking into the water, suggesting that not only the SSC motion in the trunk muscle group but also the buoyancy peculiar to underwater exercise may affect the rotation motion. Because only the propulsive phase was considered in this study, the rotational moment generated around the long axis of the trunk, which includes the recovery phase, was not taken into account. In the future, it is necessary to investigate the relationship between the factors that cause trunk rotation, including upper limb recovery motion, and the trunk muscle groups.

### Implications for the field

In short-distance crawl swimming, it was suggested that to obtain a higher swimming speed, the swimmer increases the trunk twist angle and increases the activity of the main muscles, such as the external obliques and the latissimus dorsi muscles, before the start of the main motion of the twist back motion, resulting in the SSC motion of the trunk muscle groups. This may increase the rotation angular velocity by increasing the moment exerted at the shoulder. In crawl swimming, it is important to reduce the total projected area to reduce propulsive drag, and isometric training is widely used to improve trunk muscles. However, in short-distance crawl swimming, an increase in ShRAV has been reported to be important because increased SF is a determinant of performance (Kudo et al. 2017, Koga et al. 2020). Therefore, athletes specializing in short-distance crawl swimming need to train their trunk muscles by shortening and lengthening the external obliques and the latissimus dorsi muscles, the main muscles in the trunk twisting motion. The instructor should introduce training involving relaxation and contraction in the muscle groups mobilized in the trunk twisting motion to increase not only the contraction of the muscles but also the angular velocity of the twist back, which may be important in achieving higher swimming speed in short-distance crawling.

## Conclusions

This study suggests that an increase in TA during short-distance crawl swimming induces the motion of SSC in the trunk muscle groups, which affects ShRAV in the pull-roll-back and push phases after ShR has reached its maximum value. Thus, to obtain a higher swimming speed, swimmers increase their trunk angle and increase the activity of their primary muscles, such as the external obliques and the latissimus dorsi muscles, before the start of the main motion of the twist back motion, which may induce the SSC motion in the trunk muscle group. However, we did not examine the actual muscle activity of the trunk muscles in this study, and it is necessary to investigate muscle activity during the trunk rotation motion using electromyography in the future.

## Ethics Statement

The patients/participants provided their written informed consent to participate in this study.

## Author Contributions

HH, DK, and YS contributed to the conception and design of this study. HH, DK made data acquisition. HH wrote the manuscript draft. DK, YS, and TW contributed to the manuscript revisions. All authors approved the submission of this final draft.

